# Comparative analysis of PB2 residue 627E/K/V in H5 subtypes of avian influenza viruses isolated in birds and mammals

**DOI:** 10.1101/2023.08.11.552972

**Authors:** Kelsey Briggs, Darrell R. Kapczynski

**Affiliations:** Exotic and Emerging Avian Diseases Research Unit, Southeast Poultry Research Laboratory, U.S. National Poultry Research Center, Agricultural Research Service, USDA, 934 College Station Road, Athens, GA 30605

**Keywords:** Avian Influenza virus, PB2, ANP32A, virus replication, wild birds, poultry

## Abstract

Avian influenza viruses (AIV) are naturally found in wild birds, primarily in migratory waterfowl. Although species barriers exist, many AIV have demonstrated the ability to jump from birds into mammalian species. A key contributor to this jump is the adaption of the viral RNA polymerase complex to a new host for efficient replication of its RNA genome. The AIV PB2 gene appears to be essential in this conversion as key residues have been discovered at amino acid position 627 that interact with the host cellular protein, acidic nuclear phosphoprotein 32 family member A (ANP32A). In particular, the conversion of glutamic acid (E) to a lysine (K) is frequently observed at this position following isolation in mammals. The focus of this report was to compare the distribution of PB2 627 residues from different lineages and origins of H5 AIV, determine the prevalence between historical and contemporary sequences, and investigate the ratio of amino acids in avian versus mammalian AIV sequences. Results demonstrate a low prevalence of E627K in H5 non-Goose/Guangdong/1996-lineage (Gs/GD) AIV samples, with a low number of mammalian sequences in general. In contrast, the H5-Gs/GD lineages sequences had an increased prevalence of the E627K mutation and contained more mammalian sequences. An approximate 40 percent conversion of E to K was observed in human sequences of H5 AIV, suggesting a non-exclusive requirement. Taken together, these results expand our understanding of the distribution of these residues within different subtypes of AIV and aid in our knowledge of PB2 mutations in different species.

## Introduction

Highly pathogenic (HP) avian influenza virus (AIV) outbreaks in domesticated poultry were rare prior to the 1990s [1, 2]. However, in 1996 a HPAIV H5N1 was detected in a domesticated goose (A/Goose/Guangdong/1/1996) that crossed species barriers and was detected in humans in 1997 [3, 4]. This AIV-lineage created what we now know as the H5-Gs/GD lineage and is responsible for mortality in wild birds, poultry, mammals, and humans. The H5-Gs/GD lineage viruses have become adapted and distributed across the world via migratory waterfowl. This lineage has evolved into 10 genetically distinct clades (0-9) [1]. Clade 2 versions of this lineage have become the most successful in terms of viral fitness and global distribution [1, 5, 6]. The subclade 2.3.4 was first detected in 2008 in China and has continued to evolve into the current 2.3.4.4a-h viruses [1]. The United States (U.S.) saw its first incursion of H5-Gs/GD lineage clade 2.3.4.4c viruses in 2014, which resulted in the death of over 47 million poultry resulting in an estimated loss of 3.3 billion dollars [1, 7]. Many countries, including the U.S., are experiencing large-scale outbreaks from clade 2.3.4.4b viruses, which appear very adapted to migratory waterfowl based on their global spread the past few years [8, 9].

One interest of this subclade is the adaption to other species [10-14]. A well-described avian-to-mammalian genetic adaptation is an amino acid change in the PB2 protein at residue 627. Traditionally, avian sequences contain a glutamate (E) at this position, while mammalian/human sequences contain a lysine (K) [15]. It was originally thought that this residue played a role in PB2’s ability to replicate at lower temperatures and provided an explanation for inefficient replication of AIV in non-avian hosts [16]. While temperature may still play a role, more recent structural studies have determined that PB2 interacts with host protein acidic nuclear phosphoprotein 32 family member A (ANP32A) at residue 627, and this interaction is the driving force behind the E627K mutation [17, 18]. The interaction is critical for stabilization of the AIV polymerase complex (vPol), and the amino acid composition of ANP32A that surrounds residue 627 plays a major role in supporting AIV replication [18]. However, other PB2 mutations have been found to support AIV replication in the absence of E627K, namely at positions 271, 590, and 591 [19-21].

Most avian ANP32A proteins encode an additional 33 amino acids (ANP32A_33_) in between the two domains that are critical for AIV polymerase activity [22]. The first four amino acids of the insert are a SUMO interacting motif (SIM) site that has been shown to increase binding efficiency of ANP32A_33_ with the vPol and is located directly above the PB2 627 residue [18, 23]. The SIM site contains a mixture of acidic and basic residues, which provides stabilization of the complex, and likely allows for replication with either a PB2 627E or K. The other 27 additional amino acids are duplicated from exon 4 (amino acid residues 149-175) are believed to strengthen the interaction between the vPol and ANP32A. Human, mammals, and ratite species lack the 33 amino acid insertion (ANP32A_Δ_), which results in a weaker interaction of ANP32A and the vPol leading to lower polymerase activity [17, 18, 23]. The E627K mutation appears to compensate for the weaker interaction and helps to restore vPol activity of AIV in hosts lacking ANP32A_33_ [18, 23].

Avian ANP32A contain natural variations in splicing patterns that result in three major isoforms of the protein, whereas humans only carry the ANP32A_Δ_ isotype [22, 24]. Chicken, turkey, and duck, produce all three transcripts of ANP32A (ANP32A_33_, ANP32_29_ (lacking SIM), and ANP32A_Δ_), but the ANP32A_33_ isotype is the preferentially expressed isoform (appx 65%). However, some species of wild waterfowl and other migratory birds express higher proportions of the human-like ANP32A (ANP32A_Δ_) and the partial insertion that lacks the SIM site (ANP32_29_), indicating ANP32A expression is strongly associated with host range [22-24].

In this work, we investigate the number of submitted sequences containing a PB2 627E, K or V from H5 AIV. We compare the prevalence of sequences with a 627K between non-Gs/GD and Gs/GD lineages and examine the ratio of sequences within the Gs/GD lineage. Finally, we examine residue specificity within the broad host range of clade 2 AIV, including current global 2.3.4.4 viruses.

## Methods

### Sequence analysis

All sequences were obtained from the Global Initiative on Sharing All Influenza Data (GISAID EPIFLUTM) database [25]. The search parameters on GISAID always included type A, H5, and required a complete segment of PB2. Other parameters such as host(s) and clade(s) were chosen as needed. Downloaded data sets were aligned in Geneious Prime (Boston, MA) using a MUSCLE alignment. Only complete and correct PB2 protein sequences with coverage over residue 627 were used for analysis. Sequences that did not meet this criterion were discarded. Tables were created based on the number of PB2 sequences that matched the criteria set, such as host species, total number, amino acid residue, and clade. Totals were calculated by adding up each group in the table, and in some cases the total does not represent the entire data set. Very few sequences had 627 residues that were not a glutamate (E), lysine (K), or valine (V), so they were not included in the analysis total. The percentage of K residues was obtained by dividing the number of K residues by the total of that group then multiplying by 100. When a clade or subclade was chosen in the “clade” search panel, all subclades were included in the analysis, unless otherwise noted. Tables including a species section were chicken/turkey only, all other avian species (besides chicken/turkey), mammals (not including human), and humans. It is important to note that all timespans listed in this report are based on the dataset used for analysis. They do not represent the exact circulation of those virus clades. The findings of this study are based on data from 125,996 PB2 sequences available on GISAID from March 2023.

## Results

### Prevalence of 627K in non-A/Gs/GD/96 lineage H5Nx viruses

Using the GISAID “clade” search panel, we examined sequences that were not classified as part of the Gs/Gd lineage [25]. The American-non-Gs/GD/96 (Am_nonGsGD) lineage had 789 complete PB2 sequences with 627 coverage, and the earliest viruses in the dataset were from Wisconsin, USA in 1975. The subtypes in 1975 included H5N2, H5N6, and H5N1. The species in this lineage consisted of chicken/turkey (220), all other avian (567), mammalian (2), and human (0). Of the 789 sequences examined only 2 had a PB2 627K, the animals were a rhea and emu (both Texas, USA, 1993, H5N2). The percent of PB2 627K sequences for the Am_nonGsGD lineage was 0.25%. Unexpectedly, there were 17 chicken sequences with a V at position 627, which accounted for 2.15% of the total. They originated from a H5N2 isolation in Mexico in 2019 (Table 1, top).

**Table 1:** Prevalence of PB2 627 E/K/V in non-A/Gs/GD/96 H5 lineage viruses.

| Lineage | Species | Total numbers | 627E | 627K | 627V | Percent 627E | Percent 627K |
| --- | --- | --- | --- | --- | --- | --- | --- |
| American-non-Gs/GD/96 (1975-present) | Chicken/Turkey | 220 | 203 | 0 | 17 | 92.3 | 0 |
|  | Other Avian | 567 | 564 | 2<sup>a</sup> | 0 | 99.5 | 0.4 |
|  | Mammalian | 2 | 2 | 0 | 0 | 100.0 | 0 |
|  | Human | 0 | 0 | 0 | 0 | 0 | 0 |
|  | **Total** | **789** | **769** | **2** | **17** | **97.3** | **0.25** |
| Eurasian-non-Gs/GD/96 (1959-present) | Chicken/Turkey | 46 | 0 | 0 | 0 | 0 | 0 |
|  | Other Avian | 342 | 338 | 4<sup>b</sup> | 0 | 98.8 | 1.2 |
|  | Mammalian | 2 | 2 | 0 | 0 | 100.0 | 0 |
|  | Human | 0 | 0 | 0 | 0 | 0 | 0 |
|  | **Total** | **390** | **2** | **4** | **0** | **99.4** | **1.03** |

<sup>a</sup> 1 rhea and 1 emu.
<sup>b</sup> 4 ostrich.

For the Eurasian-non-Gs/GD/96 (Ea_nonGsGD) lineage, 390 sequences were examined, the earliest sequences were from Scotland in 1959 (H5N1). In this lineage there were 46 chicken/turkey, 342 other avian, 2 mammals, and 0 human sequences. Of the 390 sequences, 4 had a PB2 627K. All 4 sequences came from ostriches in South Africa in 2011 and 2015. The percentage of PB2 627K sequences from the Ea_nonGsGD lineage was 1.03% (Table 1, bottom). Interestingly, there were only 4 mammalian sequences in the dataset within the two lineages. All 4 were H5N2 sequences from swine, two were from Mexico in 2014/2015 and two were from Korea in 2008 (Table 1). Based on the data available, the American and Eurasian lineages appear to have low mammalian/human spillover events and a low percentage of PB2 627K adaptations.

### Prevalence of PB2 627K in A/Gs/GD/96 H5 lineage clades 0-9

Next, we examined the percentage of PB2 627K residues in clades 0-9 of the Gs/GD lineage, which began in 1996. PB2 sequence availability ranged from 17-10,734 sequences between the clades (Table 2). Clade 0 had 11.7% (7/60) PB2 sequences with a 627K, which included an ostrich and 6 human samples. The dataset contained sequences from 1996-2008 (Table 2). Clade 1 had 395 sequences available, 39 sequences had a PB2 627K (9.9 %), which consisted of 14 avian species, 24 mammalian/human species, 1 unknown, and were sequenced from 1996-2014 (Table 2). Clade 2 had the most sequences available (10,734), of those 738 contained a PB2 627K and 16 contained a 627V. Clade 2 had 6.9% of total sequences with a 627K, of those 608 were avian and 130 were mammalian/human. Available clade 2 sequences ranged from 1996-present (Table 2). Clade 3 sequences contained 2.9% PB2 627K residues (6/206), 1 was from the environment and 5 were human cases. The sequences were collected from 1997-2015 (Table 2). Clade 6 (2002-2022) had 2 human cases with a PB2 627K out of 33 total, making the percentage 6.1% (Table 2). Clade 7 had 2.5% sequences with a PB2 627K, there were 2 avian samples out of 80 from the years 2002-2015 (Table 2). Clade 9 sequences ranged from 1997-2006, only 2 avian sequences had a PB2 627K out of 49 total (4.1%) (Table 2). Clades 4,5, and 8 had no sequences containing a PB2 627K (Table 2). The overall average of PB2 627K in the Gs/GD lineage was 4.9%, which is 13x more than the Am_nonGsGd and 4.8x more than the Ea_nonGsGd lineages.

**Table 2:** Prevalence of PB2 627 E/K/V in A/Gs/GD/96 H5 lineage clades 0-9.

| Clade | Total numbers | 627E | 627K | 627V | Percent 627E | Percent 627K |
| --- | --- | --- | --- | --- | --- | --- |
| 0 | 60 | 53 | 7<sup>a</sup> | 0 | 88.3 | 11.7 |
| 1 | 395 | 356 | 39<sup>b</sup> | 0 | 90.1 | 9.9 |
| 2 | 10734 | 9996 | 738<sup>c</sup> | 15 | 93.1 | 6.9 |
| 3 | 206 | 200 | 6<sup>d</sup> | 0 | 97.1 | 2.9 |
| 4 | 18 | 18 | 0 | 0 | 100.0 | 0.0 |
| 5 | 22 | 22 | 0 | 0 | 100.0 | 0.0 |
| 6 | 33 | 31 | 2<sup>e</sup> | 0 | 93.9 | 6.1 |
| 7 | 80 | 78 | 2<sup>f</sup> | 0 | 97.5 | 2.5 |
| 8 | 17 | 17 | 0 | 0 | 100.0 | 0.0 |
| 9 | 49 | 47 | 2<sup>g</sup> | 0 | 95.9 | 4.1 |

<sup>a</sup> Clade 0 – 1 avian (ostrich) & 6 human.
<sup>b</sup> Clade 1 – 14 avian (1 ostrich), 24 mammalian/human, 1 unknown.
<sup>c</sup> Clade 2 – 608 avian, 130 mammalian/human.
<sup>d</sup> Clade 3 – 1 environment, 5 human.
<sup>e</sup> Clade 6 – 2 human.
<sup>f</sup> Clade 7 – 2 avian.
<sup>g</sup> Clade 9 – 2 avian.

### Prevalence of PB2 627K in A/Gs/GD/96 H5 lineage clade 2

To examine clade 2 more thoroughly, we determined the percent PB2 627K in subclades 2.1-2.5, based on available GISAID sequences [25]. Clade 2.1 had a low proportion of avian sequences (34.6%) available compared to mammalian/human sequences (65.4%). All 19 (8.3%) sequences with a PB2 627K in clade 2.1 were mammalian/human in origin (Table 3, top). The sequences from clade 2.1 were obtained between 2003-2015. Clade 2.2 was previously shown to have a high incidence of PB2 627K residues [20]. We found that 92.1% of available PB2 sequences classified as clade 2.2 had a 627K residue. Unexpectedly, 578 sequences with a PB2 627K were from avian species and only 36 were mammalian/human. Clade 2.2 sequences ranged from 1997-2017 in the dataset (Table 3, top). Clade 2.2 contributed to 83% (614/738) of the total clade 2 PB2 627K population (Table 2). Clade 2.3 (2003-present) had the largest total number of sequences available (9,797), but only 105 had PB2 627K. Of the sequences with a 627K, 75 were mammalian/human and 30 were of avian origin. The total percentage of PB2 627K sequences for clade 2.3 was 1.1% (Table 3, top). Of note, only clade 2.3 contained PB2 627V sequences. The species containing a PB2 627V were mixed, 8 chicken, 2 other avian, 3 red foxes, 1 tiger, and 1 human. Clades 2.4 and 2.5 had low sample sizes and none of the sequences contained a PB2 627K residue. Additionally, the sequences available were only from 2003-2004 and 2003-2006, respectfully (Table 3, top). The data indicates that the majority of PB2 627K sequences are from clade 2.2, not 2.3.

**Table 3:** Comparative prevalence of PB2 E/K/V in Gs/GD/96 H5 lineage clade 2.1-2.5 (Top) and distribution within various species in subclade 2.3.4.4 (Bottom).

| Clade | Total numbers | Percent Avian Sample | 627E | 627K | 627V | Percent 627E | Percent 627K |
| --- | --- | --- | --- | --- | --- | --- | --- |
| 2.1<sup>a</sup> | 228 | 34.6 | 209 | 19<sup>d</sup> | 0 | 91.7 | 8.3 |
| 2.2<sup>b</sup> | 667 | 94.2 | 53<sup>e</sup> | 614<sup>f</sup> | 0 | 7.9 | 92.1 |
| 2.3<sup>c</sup> | 9797 | 92.4 | 9672 | 105<sup>g</sup> | 15 | 98.7 | 1.1 |
| 2.4 | 16 | 100.0 | 16 | 0 | 0 | 100.0 | 0.0 |
| 2.5 | 26 | 100.0 | 26 | 0 | 0 | 100.0 | 0.0 |
| Species |  | 2.3.4.4a (502)<sup>i</sup> | 2.3.4.4b (5311) | 2.3.4.4c (1056) | 2.3.4.4e (551) | 2.3.4.4g (84) | 2.3.4.4h (373) |
| Chicken/Turkey |  | 0 | 16 | 0 | 0 | 0 | 2 |
| Other Avian |  | 2<sup>h</sup> | 7<sup>i</sup> | 0 | 0 | 0 | 0 |
| Mammalian |  | 0 | 28<sup>j</sup> | 0 | 0 | 0 | 4<sup>k</sup> |
| Human |  | 4 | 2 | 0 | 3 | 0 | 9 |
| Percent 627E |  | 98.8 | 99.0 | 100.0 | 99.5 | 100.0 | 96.0 |
| Total with 627K |  | 6 | 53 | 0 | 3 | 0 | 15 |
| Percent 627K |  | 1.2 | 1.0 | 0.0 | 0.5 | 0.0 | 4.0 |

<sup>a</sup>2.1 includes 2.1.1, 2.1.2, 2.1.3, 2.1.3.1, 2.1.3.2, 2.1.3.2a, 2.1.3.2b, 2.1.3.3.
<sup>b</sup>2.2 includes 2.2, 2.2.1, 2.2.1.1, 2.2.1.1a, 2.2.1.2, 2.2.2, 2.2.2.1.
<sup>c</sup>2.3 includes 2.3.1, 2.3.2, 2.3.2.1, 2.3.2.1a, 2.3.2.1b, 2.3.2.1c, 2.3.3, 2.3.4, 2.3.4.1, 2.3.4.2, 2.3.4.3, 2.3.4.4, 2.3.4.4b, 2.3.4.4c, 2.3.4.4e, 2.3.4.4g, 2.3.4.4h.
<sup>d</sup>19 mammalian/human. Of mammalian/human (149), 12.8% have 627K.
<sup>e</sup>3 mammalian and 50 avian.
<sup>f</sup>36 mammalian/human and 578 avian (4 ostrich).
<sup>g</sup>75 mammalian/human and 30 avian (1 emu, 2 rhea).
<sup>h</sup>1 cormorant and 1 pheasant.
<sup>i</sup>2 rhea, 1 emu, 1 duck, 2 common tern, and 1 unknown avian.
<sup>j</sup>1 bear, 2 skunk, 3 cat, 2 ferret, 1 racoon, 14 foxes, 4 seals, and 1 otter.
<sup>k</sup>4 mink.
<sup>i</sup> number of samples per subclade.

The most current and prevalent Gs/GD lineage viruses belong to clade 2.3.4.4 [1]. We wanted to examine which species in subclades 2.3.4.4a-h had a PB2 627K residue. The subclades 2.3.4.4d and 2.3.4.4f were not “clade” options in the GISAID database, so they were not included in the analysis [25]. Clade 2.3.4.4a contained 1.2% (6/502) sequences with a PB2 627K, the species included 2 avians and 4 humans. Sequences in this clade were collected between 2012 and 2018 (Table 3, bottom). The 2.3.4.4b viruses are responsible for the current outbreak in the U.S. and have been reported in many countries [12, 25]. Of the 5,311 sequences analyzed from years 2003-present, 53 had a PB2 627K making the percentage only 1.0%. However, 48 of the 53 sequences were from 2021 to present. Bird sequences containing a 627K accounted for 23 of the 53 sequences and included chicken/turkeys, ducks, ratites, and common terns. There was a higher diversity of mammalian species containing a PB2 627K in the 2.3.4.4b viruses, compared to all other groups analyzed. Most of the mammalian/human sequences in clades 2.1 and 2.2 were human, whereas the 2.3.4.4b sequences included a bear, 2 skunk, 3 cats, 2 ferrets, 1 raccoon, 14 foxes, 4 seals, 1 otter, and 2 human (Table 2 and Table 3, bottom). Of note, 2.3.4.4c sequences, which were responsible for the first outbreak of a Gs/GD lineage H5Nx virus in the U.S., did not have any sequences with PB2 627K [8]. The dataset contained 1,056 samples, and included sequences from 2014-2020 (Table 3, bottom). Three human samples (3/551) from 2.3.4.4e had PB2 627K from years 2014-2018 (Table 3, bottom). Clade 2.3.4.4g (2014-2020) had the smallest sample size (84), and none of the sequences contained PB2 627K (Table 3, bottom). Lastly, 2.3.4.4h had the highest percentage of PB2 627K sequences at 4.02% (15/373), spanning from 2014 to 2021. The sequences were from 2 chicken/turkey, 4 mink, and 9 humans (Table 3, bottom). Taken together, the data indicates that the current 2.3.4.4b viruses have a low overall percentage of PB2 627K sequences, but an increased propensity to infect multiple species.

### Prevalence of PB2 627E/K/V in human sequences from different hemagglutinin subtypes

Finally, we examined the prevalence of PB2 627K in human AIV sequences with more common HA subtypes [26]. First, the percent of PB2 627K in human sequences in subtypes (H1-H3) was examined. Interestingly, 95% of H1 sequences sequenced contained the avian-like PB2 627E. We only observed 4.7% of all H1 sequences demonstrating the PB2 627K residue. However, the majority of the 39,418 sequences examined were from the pandemic H1N1 (pdm09) lineage, which began in 2009, and was known to contain a PB2 segment from an avian North American virus (Table 4) [27]. As expected, both the H2 and H3 sequences contained higher percentages of PB2 627K residues with prevalence at 91.9% and 99.4%, respectfully (Table 4). Next, we examined the proportion of PB2 627K in avian-adapted AIV subtypes (H5, H7, and H9). H5 viruses obtained from human samples contained 39.1% PB2 627K sequences (Table 4). Of the 1,207 human sequences classified as subtype H7, 851 or 70.4% contained a K at position 627 (Table 4). Lastly, the H9 subtype contained only 2 human sequences with a PB2 627K (Table 4). Interestingly, the H7 and H9 viruses contained a higher proportion of 627V residues, 2.7% and 36%, respectively. Apart from the pdm09 H1N1 viruses, the data illustrates that human-adapted viruses typically contain a PB2 627K.

**Table 4:** Prevalence of PB2 627 E/K/V from human sequences amongst different influenza hemagglutinin subtypes.

| HxNx | Total numbers | 627E | 627K | 627V | Percent 627E | Percent 627K |
| --- | --- | --- | --- | --- | --- | --- |
| H1 | 39418 | 37559 | 1855 | 4 | 95.3 | 4.7<sup>a</sup> |
| H2 | 123 | 10 | 113 | 0 | 8.1 | 91.9 |
| H3 | 71994 | 419 | 71560 | 15 | 0.6 | 99.4 |
| H5 | 402 | 244 | 157 | 1 | 60.7 | 39.1 |
| H7 | 1207 | 323 | 851 | 33 | 26.8 | 70.4<sup>b</sup> |
| H9 | 59 | 36 | 2 | 21 | 61.0 | 3.4 |

<sup>a</sup>Pandemic H1N1 (pdm09) contained a PB2 from an avian North American virus.
<sup>b</sup>Recent H7N9 cases predominate sequences in this subtype.

## Discussion

The goal of this research was to compare the distribution of PB2 627 residues between avian and mammalian sequences in H5 from non-Gs/GD and Gs/GD lineages and determine the prevalence between previous subclades and current ones. The current clade 2.3.4.4b H5Nx AIVs have a global distribution and raise concerns about mammalian adaptions as recent isolations have occurred in domestic and peridomestic mammals [12, 28]. Here, we focused on one well known avian to mammalian adaption residue, E627K/V, to determine if the 2.3.4.4 viruses have an increased propensity to mutate in that direction.

All sequences were obtained from GISAID, so the number of sequences were limited to what was available in the database. We chose GISAID because it contained the most sequences of current (2021-present) AIV, but it is possible that older strains were missed in the analysis because they were not added retroactively (prior to 2008) [25]. AIV surveillance and reporting isn’t a standard practice in all countries, consequently samples are limited to countries who do report, and this may contribute to the low dataset numbers and sample bias in some clades. Obtaining samples from only dead birds may also result in sampling bias, but it’s not possible to tell whether samples were taken during active or passive surveillance.

In the non-Gs/GD lineage, the majority of sequences contained the 627E residue. We observed only 6 avian sequences contained a PB2 627K and they belonged to the Ratite family (Ostrich, Rhea, Emu, and Cassowary and Kiwi) of birds (Table 1). Ratites were the only species of birds that contained PB2 627K residues in this group. Phylogenetic analysis of the ANP32A gene demonstrates that the Ratite family lack the 33 amino acid insertion in exon 5 that most other avian species have, which presumably make it more mammalian like, and may explain why viruses isolated from these species select for 627K during replication [14, 22]. Of further note is that both rhea and emu sequences from the Am_nonGsGD lineage contained multiple polybasic residues at the HA cleavage site, suggesting they were HPAI viruses. In the EA_nonGsGD lineage, the two ostriches from 2011 contained a HPAI cleavage sequence, whereas the two ostriches from 2015 contained a LPAI sequence.

Clade 2 is the most evolutionarily successful clade from the Gs/GD lineage of AIV based on sample size and the number of subclades (Table 2 & 3) [1, 29]. Within subclade 2, the proportion of avian and mammalian samples in clade 2.1 was unexpected. The largest portion of samples were human sequences from Indonesia ranging from 2003 to 2015 (Table 3). Of note is the observation that most of the sequences in clade 2.2 containing a 627K were of avian origin (Table 3). Clade 2.2 viruses were shown to be shed via the respiratory route in waterfowl rather than the typical cloacal route, therefore it was proposed that clade 2.2 viruses maintained the 627K to compensate for the cooler temperature of respiratory tract [20, 30, 31].

Recently, a study using RNA-seq showed that some waterfowl, land fowl, and pelicans preferentially express a human-like ANP32A, which also could have contributed to a larger proportion of avian species containing a 627K in clade 2.2, as there was no selective pressure to maintain 627E [22]. Additionally, Long et al. found that the PB2 627K mutation did not affect pathogenesis or transmission in ducks suggesting that the mammalian adaption could be maintained in an avian species, which also supports the notion that there is little selection pressure to go from K627E [20]. The clade 2.3 dataset was comprised of 17 subclades, including the current 2.3.4.4b viruses. Interestingly, 98.7% (9672) of the 2.3 sequences contained a 627E residue despite only 92.4% (9052) being of avian origin, demonstrating that non-avian species also maintained the 627E residue. This observation went both ways in that of the 105 clade 2.3 sequences demonstrating a 627K, only 71.4% (75) were mammalian (Table 3A).

Within clade 2.3.4, more than half of the PB2 627K sequences were in the clade 2.3.4.4b subclade (Table 2 and 3). There were 16 chicken/turkey species and 7 other avians that contained a PB2 627K in clade 2.3.4.4b, this may indicate that the virus is spilling over from mammals into avians and maintaining the residue at the time of isolation. Several accounts of mammalian spillover events have occurred since the 2.3.4.4b viruses have become predominant, and the diversity of species being infected is unprecedented (Table 3) [9, 12, 13, 32]. Nevertheless, based on our data, the number of 2.3.4.4b mammalian sequences with a PB2 627K is still lower than the sequences with a 627E. Interestingly, we found that 5 sea mammals (4 seals and 1 otter) contained a PB2 627K, whereas a recent study examined dolphin and sea lion sequences from Peru and found they contained a 627E. Leguia et al. also proposed that transmission between sea lions off the coast of Peru could be occurring rather than independent avian spillover events because of the massive die-offs being observed [28]. More investigation is required to determine if mammal-to-mammal and mammal-to-avian transmission is occurring, and which residues are allowing for efficient replication of the virus.

Despite the fact that residue PB2 627 is almost exclusively a K or E in all influenza A viruses, a small portion contain a valine (V). A study by Chin et al. found inducing random mutations at position 627 allowed for the 627V mutation when purified in a mammalian system but not in an avian system [33]. This may indicate that the 627V mutations observed in all avian species were transmitted from a mammalian host (Tables 1-4). The study observed a slight reduction in replication compared to a 627K mutation using the culture-adapted A/Puerto Rico/8/34 (H1N1) virus [33]. However, a more recent study used the polymerase genes from an avian H5N1 (A/Muscovy duck/Vietnam/TY93/2007; clade 2.3.4) to rescue a virus containing the PB2 627V mutation. Taft et al. found that the 627V mutation had significantly increased viral replication in mammalian cell culture and virulence in mice that was comparable to 627K virus [34]. Additionally, Luk et al. found that an H7N9 virus containing a PB2 627V was extremely fit and transmissible between chickens and mammals [35]. The number of V residues in H5 sequences remain low, however the H7 and H9 human cases had a considerable number of sequences containing a 627V, many of which are from recent years (Table 4).

The dependency of viral polymerases on host ANP32A is one factor to cross host barriers, exemplified by the propensity for AIV sequences to mutate to a 627K in mammalian sequences (Table 1-4) [36]. The Gs/GD lineage viruses are commonly found in wild waterfowl that appear to transcribe higher rates of the human-like ANP32A, which may account for shifts in 627 residue specificity [36, 37]. While PB2 627K is an established marker for mammalian adaption, it is not solely responsible for it [15, 27, 33]. The pdm09 H1N1 viruses contained a PB2 627E, due to the avian origin of the segment but had other compensatory mutations that allowed for efficient replication in mammals and humans [19, 20, 38]. As more mammalian species become infected by 2.3.4.4b AIV additional residues may yet be identified. It is known that efficient replication in a certain host requires adaption to the machinery within, and that adaption from one host species to another requires mutation and selection. In this work, we performed a differential analysis of PB2 residue 627 and demonstrate a non-exclusive requirement for conversion.

## Acknowledgements

We thank Ryan Sweeney for excellent technical assistance. We gratefully acknowledge all data contributors, i.e., the Authors and their Originating laboratories responsible for obtaining the specimens, and their Submitting laboratories for generating the genetic sequence and metadata and sharing via the GISAID Initiative, on which this research is based.

## Author contributions

**Kelsey Briggs:** Conceptualization, Methodology, Formal analysis, Writing – original draft, preparation, Writing – review & editing. **Darrell R. Kapczynski:** Conceptualization, Methodology, Formal analysis, Writing – original draft, preparation, Writing – review & editing.

## Data availability

The data are available from the senior author upon request.

